# Testing SIPA1L2 as a modifier of CMT1A using mouse models

**DOI:** 10.1101/2023.11.30.569428

**Authors:** George C. Murray, Timothy J. Hines, Abigail L.D. Tadenev, Isaac Xu, Stephan Züchner, Robert W. Burgess

**Affiliations:** The Jackson Laboratory, Bar Harbor, ME 04609; The Graduate School of Biomedical Science and Engineering, The University of Maine, Orono, ME 04469; Department of Human Genetics and John P. Hussman Institute for Human Genomics, University of Miami Miller School of Medicine, Miami, FL, USA

## Abstract

Charcot-Marie-Tooth 1A is a demyelinating peripheral neuropathy caused by the duplication of peripheral myelin protein 22 (*PMP22*), which produces muscle weakness and loss of sensation in the hands and feet. A recent case-only genome wide association study by the Inherited Neuropathy Consortium identified a strong association between variants in signal induced proliferation associated 1 like 2 (*SIPA1L2*) and strength of foot dorsiflexion. To validate *SIPA1L2* as a candidate modifier, and to assess its potential as a therapeutic target, we engineered mice with a deletion in *SIPA1L2* and crossed them to the C3-PMP22 mouse model of CMT1A. We performed neuromuscular phenotyping and identified an interaction between *Sipa1l2* deletion and muscular endurance decrements assayed by wire-hang duration in C3-PMP22 mice, as well as several interactions in femoral nerve axon morphometrics such as myelin thickness. Gene expression changes suggested an involvement of *Sipa1l2* in cholesterol biosynthesis, which was also implicated in C3-PMP22 mice. Though several interactions between *Sipa1l2* deletion and CMT1A-associated phenotypes were identified, validating a genetic interaction, the overall effect on neuropathy was small.

## INTRODUCTION

Charcot-Marie-Tooth disease (CMT) is a collection of inherited peripheral neuropathies that result in demyelination and axon degeneration in motor and sensory axons of the peripheral nervous system (SAPORTA AND SHY 2013). Although almost 100 loci have been associated with CMT in the human genome, CMT1A is by far the most common form, accounting for at least a third of all cases (DIVINCENZO *et al*. 2014). CMT1A is caused by unequal crossover at repeat sequences that flank 1.5 megabases of DNA on chromosome 17p12, resulting in a duplication of the intervening sequence (LUPSKI *et al*. 1991; LUPSKI *et al*. 1992; PATEL *et al*. 1992). This duplication includes the peripheral myelin protein 22 (*PMP22*) gene, and the increased gene dosage of *PMP22* underlies the demyelinating neuropathy of CMT1A.

Although CMT is commonly considered a monogenic (Mendelian) disease, genetic and environmental modifiers can influence the clinical presentation and severity. This is particularly evident in CMT1A, where the patient duplications are quite homogeneous, but clinical signs are still variable. A recent case-only GWAS study performed by the Inherited Neuropathy Consortium (INC) identified four suggestive loci that associated with outcomes including difficulty in eating with utensils, hearing loss, decreased ability to feel, and the CMT neuropathy score (TAO *et al*. 2019a). However, the strongest association was between weakness in foot dorsiflexion with four SNPs in the *SIPA1L2* gene (TAO *et al*. 2019b). Interestingly, this more comprehensive genome-wide analysis did not find associations for two previously implicated modifiers, *LITAF*, which can cause CMT on its own, and miRNA 149 (MEGGOUH *et al*. 2005; SINKIEWICZ-DAROL *et al*. 2015; NAM *et al*. 2018).

SIPA1L2 contains a RAP/RAN GTPase activating domain (GAP), a PDZ domain, and a C-terminal coiled-coil domain. It associates with postsynaptic density proteins and is enriched at synapses in cultured hippocampal neurons, but unlike other SIPA family members, over expression does not alter synaptic spine morphology (SPILKER *et al*. 2008). It also associates with myosin heavy chain 9 and beta-actin (TAO *et al*. 2019b). SIPA1L2 has been shown to regulate retrograde trafficking of BDNF/TrkB amphisomes, and *Sipa1l2* knockout mice show impaired BDNF-dependent presynaptic plasticity (ANDRES-ALONSO *et al*. 2019). Importantly for myelination, it is expressed in Schwann cells and is regulated by SOX10, putting it in a co-expression network with other myelin genes. Knockdown of *Sox10* results in lower *Sipa1l2* levels, and knockdown of *Sipa1l2* results in decreases in other relevant genes including *Pmp22*, *Mpz*, and *Egr2*, suggesting *Sipa1l2* is in the same SOX10 co-expression network as other myelin genes (TAO *et al*. 2019b). In addition to the recent association with CMT1A, it is also associated with Parkinson’s disease in multiple (but not all) patient cohorts (CHEN *et al*. 2016; SAFARALIZADEH *et al*. 2016; WANG *et al*. 2016; FOO *et al*. 2017; YANG *et al*. 2017; ZOU *et al*. 2018).

The GWAS findings suggest that *SIPA1L2* may be a modifier of CMT1A (TAO *et al*. 2019b). They also suggest that *SIPA1L2* may be a therapeutic target for CMT1A, based on the idea that reducing *SIPA1L2* expression may concomitantly decrease the expression of other myelin genes including *PMP22*. The GWAS association of *SIPA1L2* with foot dorsiflexion and CMT1A is impressive for the size of the study but is still below the normal statistical threshold for genome-wide significance and therefore requires additional validation. Rare disease GWAS are very challenging because of low patient numbers, making additional studies in patients impractical. Finally, the effect of *SIPA1L2* on myelin gene expression was identified *in vitro* in cell lines (TAO *et al*. 2019b). Reproducing the effect *in vivo* would strongly support *SIPA1L2*’s involvement in the co-expression network.

We first searched sequencing data from 50 CMT1A patients to look for up-/ down-stream variation in the vicinity of the *SIPA1L2* gene that could impact gene expression, focusing on haplotypes that contained the *SIPA1L2* SNPs reported in the original GWAS study. No additional coding variants, variants in canonical splice junctions, or structural changes were detected. We decided that, in mice, a larger deletion in *Sipa1l2* would provide better insight into gene function and disease-relevant interactions than replicating intronic SNPs from the GWAS. We used CRISPR/Cas9 genome editing to delete 1,877bp at the 5’-end of the *Sipa1l2* gene. We also obtained the C3-*PMP22* transgenic mouse model of CMT1A (VERHAMME *et al*. 2011). These mice carry a human YAC transgene containing the *PMP22* gene, and thus accurately model the increased gene dosage of CMT1A. They develop a relevant and progressive demyelinating phenotype (VERHAMME *et al*. 2011). Mice with the *Sipa1l2* deletion were bred to the C3-*PMP22* transgenic mice to assess the effects of altering *Sipa1l2* levels on the severity of the demyelinating phenotype of C3-*PMP22*. We performed an analysis of neuromuscular phenotypes, including neurophysiology and histopathology. We also performed RNAseq to assess changes in myelin gene expression and to identify altered or interacting pathways.

Here we show that *Sipa1l2* deletion does not induce neuromuscular phenotypes on its own. We identify interactions between the *Sipa1l2* deletion and C3-PMP22 phenotypes such as muscular endurance, body weight, and myelin thickness. Our gene expression analysis identifies a potential role for *Sipa1l2* in cholesterol biosynthesis and an interaction with C3-PMP22 gene expression signatures. However, we failed to detect changes in SOX10/EGR2 network genes at 6mo of age, suggesting that *Sipa1l2* may interact with cholesterol biosynthesis independently of *Egr2* and other myelination related genes, or alternatively that repression of cholesterol biosynthesis is more persistent than any expression changes directly related to myelination. Taken together, our *in vivo* findings support *Sipa1l2* as a modifier of CMT1A severity. However, the interaction does not appear to markedly rescue CMT1A phenotypes in this model.

## METHODS

### Sequence analysis of human **SIPA1L2** locus

CMT1A duplication diagnosis was obtained by clinical genetic testing. Whole genome sequencing was performed from peripheral blood using the Illumina TruSeq DNA library preparation kit. DNA was sequenced to a mean depth of 30x. The generated sequencing data were then aligned to the GRCh37 reference genome using the Burrows-Wheeler aligner (version: 0.7.12) and variants were called using GATK (version 4.1.4.1) and then imported to the GENESIS genome database and analysis platform (GONZALEZ *et al*. 2015). Additional inferences with functional elements and haplotypes were made using the UCSC genome browser.

### Experimental Mice: C3-PMP22 and Sipa1l2 knockout

The C3-PMP22 mouse model of CMT1A was obtained from Dr. Frank Baas of Amsterdam University. The official strain designation of these mice is B6.Cg-Tg(PMP22)C3Fbas/J, MGI:5817396, but they will be referred to as C3-PMP22 for brevity. These mice carry a human YAC containing the PMP22 gene, and they are a derivative of the C22-PMP22 line (which carries 7-8 copies of the transgene) and have undergone a spontaneous reduction of transgene copy number (now 3-4 copies) (HUXLEY *et al*. 1996; VERHAMME *et al*. 2011). To create a *Sipa1l2* loss-of-function allele, we used CRISPR/Cas9 genome editing, employing two guides flanking the start ATG-containing exon of mouse *Sipa1l2* (guide sequences: upstream sgRNA1: TTAAAGAGTGCTGCCGAAGC, downstream sgRNA1: CTATTAGACTGACAAAGCGT). The mouse gene consists of 23 exons. The region we targeted includes 1,723 base pairs containing 5’ UTR sequence, the start codon, and the first 1,486 base pairs of coding sequence in exon 1. Two double strand breaks were repaired with the deletion of the intervening 1,877 base pairs including the entire exon sequence. These mice were bred for three generations to eliminate mosaicism and reduce possible off target mutations. C3-PMP22 transgenic mice were crossed with *Sipa1l2+/-* heterozygotes to produce *Sipa1l2+/-* mice carrying the *PMP22* transgene. These C3-PMP22::Sipa1l2+/-mice were then crossed to WT::Sipa1l2+/-mice to produce experimental genotypes: WT::WT, WT::Sipa1l2+/-, WT::Sipa1l2-/-, C3-PMP22::WT, C3-PMP22::Sipa1l2+/-, C3-PMP22::Sipa1l2-/-. Experimental cohorts were aged to 4mo and 6mo for experiments. Mice were housed under standard conditions with 14:10-hour light / dark cycles and *ad libitum* access to food and water. All procedures were performed in accordance with The Guide on the Care and Use of Laboratory Animals and were approved by the Institutional Animal Care and Use Committee of The Jackson Laboratory.

### Genotyping

Toe biopsies were obtained from mice during the first postnatal week or ear notches were collected > 14 days of age. Samples were digested in a 50 mM NaOH solution, which after neutralization with Tris buffer were then used for PCR reactions. Products were visualized using standard electrophoretic techniques to detect the C3-PMP22 transgene or the Sipa1l2 deletion. Primers are as follows: C3-PMP22 forward = CCC CTT TTC CTT CAC TCC TC and reverse = CCA ATA AGC GTT TCC AGC TC; Sipa1l2 knockout forward = CAG CCT TGC ACA ACA GGA TA, reverse = CTC CCA TCT GTG CAG CTA TCA; Sipa1l2 wildtype forward = CAG CCT TGC ACA ACA GGA TA, reverse = TGG AAG GGT CCT TGA GTT TG.

### Neuromuscular Phenotyping

Muscular endurance was evaluated using the wire hang duration test. Briefly, mice were placed on a wire mesh grid, which was then inverted, and the time they can remain suspended was recorded. Trials lasted up to one minute, and the average of three trials is reported (MOTLEY *et al*. 2011). Sciatic motor nerve conduction velocity (NCV) was measured using electrode placement described previously (MOTLEY *et al*. 2011; BURGESS *et al*. 2016; MORELLI *et al*. 2017). Mice were anesthetized with isoflurane and placed on a heating pad to maintain normal body temperature. Recording electrodes were placed in the left rear paw, and stimulating electrodes were placed near the ankle and then near the sciatic notch. NCV was calculated in m/s by dividing the distance between stimulation points by the difference in latency between the greatest amplitude of sciatic notch and ankle EMG waveforms. Bodyweights were recorded. Mice were euthanized using CO_2_ and tissues were collected. Triceps surae muscles were collected, and weights were recorded for muscle-weight-to-body-weight measurements to assess atrophy. All statistical comparisons were performed using GraphPad PRISM 9. Two-way ANOVA were applied to test differences between all genotype means, and either Tukey’s multiple comparisons test (wire-hang duration, nerve conduction velocity, muscle-to-bodyweight ratio) or Dunnett’s multiple comparisons test (bodyweight) were applied.

### Histology

Motor and sensory branches of femoral nerves were dissected free and fixed overnight in 2% glutaraldehyde and 2% paraformaldehyde in a 0.1 M cacodylate buffer. Nerves were processed for histology by Electron Microscopy Services at The Jackson Laboratory. Sections were stained with toluidine blue and imaged using a Nikon Eclipse E600 microscope with a 40X objective. Automated quantification was performed in ImageJ using threshold function and the Analyze Particles plugin as described previously (BOGDANIK *et al*. 2013). The line tool in ImageJ was used to record axon diameter and axon diameter plus myelin for 50 randomly selected axons per nerve section to calculate G-ratio (axon diameter / axon + myelin diameter). Images were visually inspected, and the number of totally demyelinated axons was recorded. Statistical comparisons were performed using GraphPad PRISM 9. Two-way ANOVA were used to test differences between all means with either Dunnett’s multiple comparisons test (axon counts) or Tukey’s multiple comparisons test (percent demyelinated and G-ratios).

### RNASeq

Tissues were lysed and homogenized in TRIzol Reagent (ThermoFisher) using a Pellet Pestle Motor (Kimbal), then RNA was isolated using the miRNeasy Micro kit (Qiagen), according to manufacturers’ protocols, including the optional DNase digest step. RNA concentration and quality were assessed using the Nanodrop 2000/Nanodrop 8000 spectrophotometer (Thermo Scientific) and the RNA 6000 Nano and Pico Assay (Agilent Technologies). Libraries were constructed using the KAPA mRNA HyperPrep Kit (Roche Sequencing and Life Science), according to the manufacturer’s protocol. Briefly, the protocol entails isolation of polyA containing mRNA using oligo-dT magnetic beads, RNA fragmentation, first and second strand cDNA synthesis, ligation of Illumina-specific adapters containing a unique barcode sequence for each library, and PCR amplification. The quality and concentration of the libraries were assessed using the D5000 ScreenTape (Agilent Technologies) and Qubit dsDNA HS Assay (ThermoFisher), respectively, according to the manufacturer’s instructions. Libraries were sequenced 150 bp paired-end on an Illumina NovaSeq 6000 using the S4 Reagent Kit v1.5. Analysis of RNASeq data was performed using an open source Nextflow pipeline (v3.5) at The Jackson Laboratory comprising tools to perform read quality assessment, alignment, and quantification (EWELS *et al*. 2020). The pipeline takes as input a single sample and outputs read counts. FastQC (RRID:SCR_014583) was used for quality checks and then Trim Galore! (RRID:SCR_011847) was used to remove adapters and sequences with low quality (Fred < 20). Sequence reads that passed quality control were aligned to mouse reference (GRCm38) using STAR (v2.7) (RRID:SCR_004463) and gene expression estimates were made using RSEM (v1.3) (RRID:SCR_013027) with default parameters. Differential expression analysis was performed using a differential expression script in R developed by Computational Sciences at The Jackson Laboratory based on the EdgeR package (ROBINSON *et al*. 2010). We selected FDR < 0.05 and absolute Log_2_FC>1.5 cutoffs to determine differential expression. We normalized raw counts values using trimmed mean of m-values (TMM) at this stage for follow-up Gene Set Enrichment Analysis.

### Over-representation and Gene Set Enrichment Analyses

ENSEMBL identifiers for differentially expressed genes were uploaded to MouseMine.org to perform over-representation analyses through: Reactome pathways, Mammalian Phenotype Ontology, and Gene Ontology (MOTENKO *et al*. 2015). We recognized at the outset that the relative paucity of differentially expressed genes would temper any conclusions drawn from this analysis. We only reported annotations with a Holm-Bonferroni corrected p-value < 0.05. To improve our ability to detect changes in pathway regulation driven by coordinated low magnitude changes across multiple gene sets, we performed Gene Set Enrichment Analysis (GSEA) using TMM counts, an approach that would be less susceptible to bias due to arbitrary cutoffs. We used default GSEA settings, which we determined were appropriate for our dataset, including a weighted enrichment statistic, the Signal2Noise method for ranking genes, and 1000 permutations. We queried against the MSigDB Reactome pathway gene lists. After a comprehensive analysis of parameterizations, we determined that an Experimental_versus_Control design focused on negative normalized enrichment scores was most informative to address our experimental question related to the interaction of *Sipa1l2* deletion with CMT1A phenotypes. We performed a GSEA leading edge analysis to identify the core driver genes associated with cholesterol biosynthesis pathways for all genotypes (SUBRAMANIAN *et al*. 2005). Each gene list was analyzed against the ENCODE database through the Network Analyst web tool to identify upstream regulators (ZHOU *et al*. 2019).

## RESULTS

### Sequence analysis of the human **SIPA1L2** locus reveals no additional variants

We performed whole genome sequencing on 50 patients with CMT1A. The goal was to evaluate the presence of variation in the 117kb genomic space of the *SIPA1L2* gene that could impact gene expression. The coding and non-coding sequence of 50 CMT1A patients was studied including 2kb upstream and downstream of the SIPA1L2 gene (NM_020808). Special attention was given to the haplotypes that contained the associated SNP (rs10910527, rs7536385, rs4649265, rs1547740) reported in the original GWAS (TAO *et al*. 2019b). No coding variation with CADD scores >18, MAVERICK>0.1 were identified (DANZI *et al*. 2023). No variation that in the canonical splice junctions or predicted to have splicing effects by SpliceAI was identified. No structural changes were present in these CMT1A samples. While this doesn’t present an exhaustive screen, it remains unclear whether the associated marker SNPs in *SIPA1L2* are correlated with pathogenic or quantitative trait loci in the vicinity of *SIPA1L2*. More comprehensive future studies, including long read sequencing will potentially clarify the presence of functionally active DNA variation.

### Mice with a CRISPR Sipa1l2 deletion do not exhibit neuromuscular phenotypes

We used CRISPR/Cas9 with two guides flanking exon 1 of the *Sipa1l2* gene on mouse chromosome 8. This produced a 1,877 bp deletion including the first 1,486 base pairs of coding sequence **(Figure 1A)**. These mice were bred for at least three generations to eliminate mosaicism and reduce possible off target mutations. We verified the deletion in homozygous *Sipa1l2-/-* mice by PCR of genomic DNA, which produced PCR products of the anticipated size **(Figure 1B)**. The 1,877 bp deletion was also verified with Sanger sequencing of PCR products spanning the deleted region in genomic DNA **(Figure 1C)**. We performed RT-PCR using lung and brain mRNA from *Sipa1l2-/-* mice and found *Sipa1l2* is not detected in homozygous knockout mice when using a primer combination that includes the coding sequence deleted in exon 1 **(Figure 1D)**. RNASeq read alignment showed that, indeed, exon 1 of *Sipa1l2* lacks coverage in homozygous knockouts; however, exons downstream of the first exon have coverage at approximately wildtype levels **(Figure 1E).**

**Figure 1.**
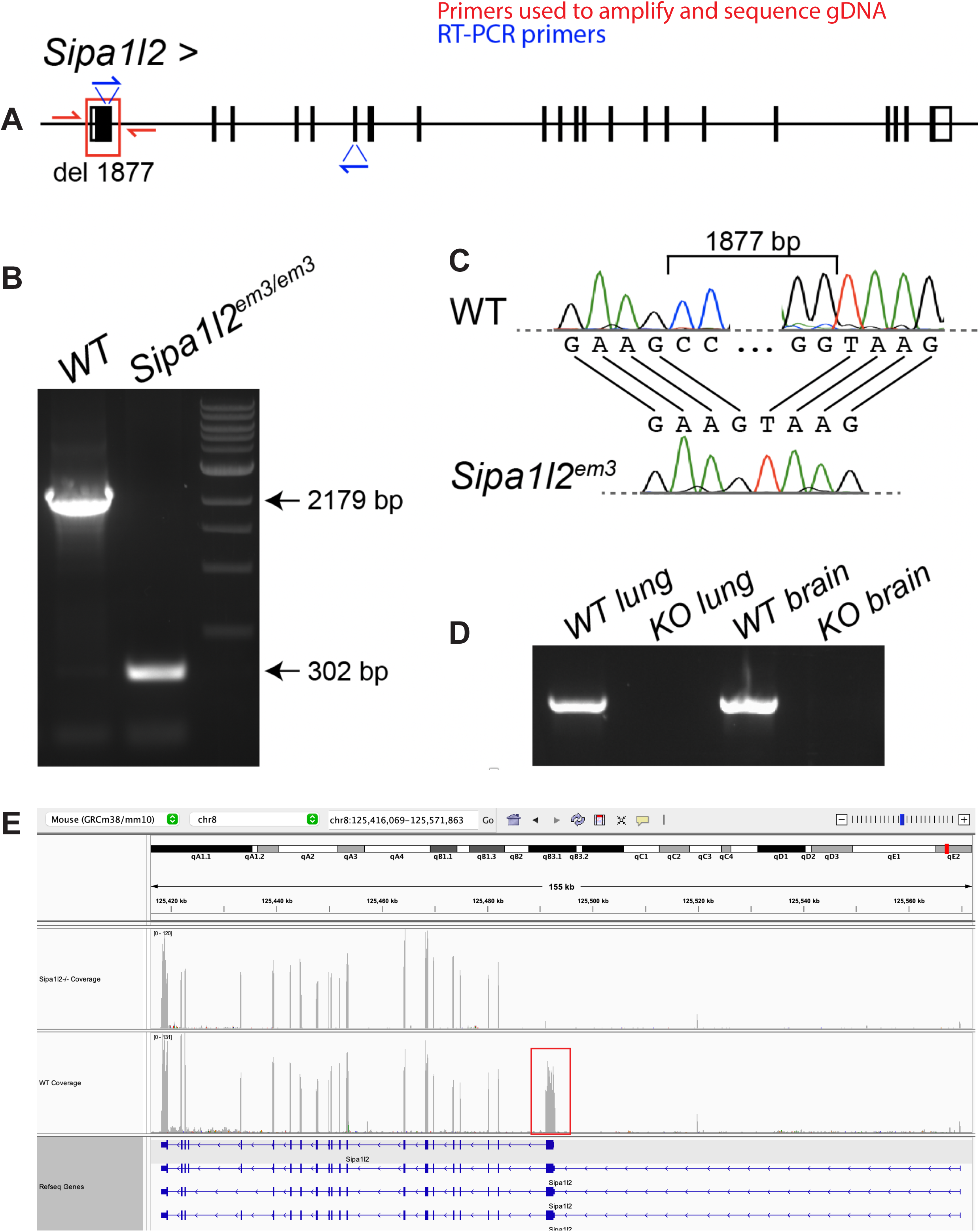
**– CRISPR engineered deletion of 1,877 bp at the 5’ end of**Sipa1l2. A) CRISPR/Cas9 genome editing was used to introduce an 1,877 bp deletion containing the first 1,486 coding nucleotides of *Sipa1l2*. The gene model is read left-to-right, inverted from its original orientation on the reverse strand for readability. The red box indicates the deletion and red half arrows indicate the position of primers used to sequence and amplify genomic DNA. Blue half arrows indicate primers used for RT-PCR. B) Gel electrophoresis of PCR products from genomic DNA showing deletion of 1,877 bp in homozygous *Sipa1l2* knockout mice. C) Sanger sequence chromatogram of genomic DNA defining the break points of the deletion in homozygous *Sipa1l2* knockout mice. D) RT-PCR from lung and brain mRNA indicates that Sipa1l2 amplicons are absent in homozygous Sipa1l2 knockout mice using a primer in the deleted region. E) RNASeq alignment in IGV from representative *Sipa1l2* knockout and wildtype control mice reveals an absence of read coverage along exon 1 of *Sipa1l2* in knockout mice. The normal wildtype coverage is indicated by the red box. However, expression was detected in downstream exons at levels similar to wild type. The gene model in this IGV view is read right-to-left reflecting that *Sipa1l2* is on the reverse strand.

### Neuromuscular phenotyping at 4mo and 6mo revealed interactions with CMT1A phenotypes

Grip strength and endurance were assayed using the wire-hang test (see Methods). Testing at 4mo identified significantly (p=0.0007) reduced wire-hang duration in C3-PMP22 mice compared to littermate wild types. C3-PMP22 mice with either heterozygous or homozygous *Sipa1l2* deletions did not differ significantly from littermate wild types, but the wire-hang duration was not significantly greater than C3-PMP22 **(Figure 2A),** indicating an intermediate phenotype. At 4mo, neither expression of the C3-PMP22 transgene nor *Sipa1l2* deletion influenced overall bodyweight **(Figure 2B)**. Sciatic nerve conduction velocity was significantly reduced in all genotypes carrying the C3-PMP22 transgene compared to wild type littermates. Wild type vs C3-PMP22 (p<0.0001), wild type vs C3-PMP22::Sipa1l2+/-(p<0.0001), and C3-PMP22::Sipa1l2-/-(p=0.0003). The *Sipa1l2* deletion did not modify conduction velocity **(Figure 2C)**. Similar tests were performed at 6mo of age. Wire-hang duration was significantly decreased in C3-PMP22 (p=0.0076) and C3-PMP22::Sipa1l2+/-genotypes (p=0.0086) compared to wild type littermates, while the C3-PMP22::Sipa1l2-/-mice were again not significantly different than control, suggesting some persistent benefit of homozygous Sipa1l2-/-deletion in the C3-PMP22 background **(Figure 2D)**. Bodyweight was significantly (p=0.012) decreased only in the C3-PMP22::Sipa1l2-/-genotype compared to wild-type littermates **(Figure 2E)**. Sciatic nerve conduction velocity was diminished (p<0.0001) in all genotypes containing the C3-PMP22 transgene, and Sipa1l2 deletion genotype did not modify nerve conduction velocity **(Figure 2F)**. Triceps surae muscle-to-bodyweight ratio did not differ significantly between any genotype.

**Figure 2.**
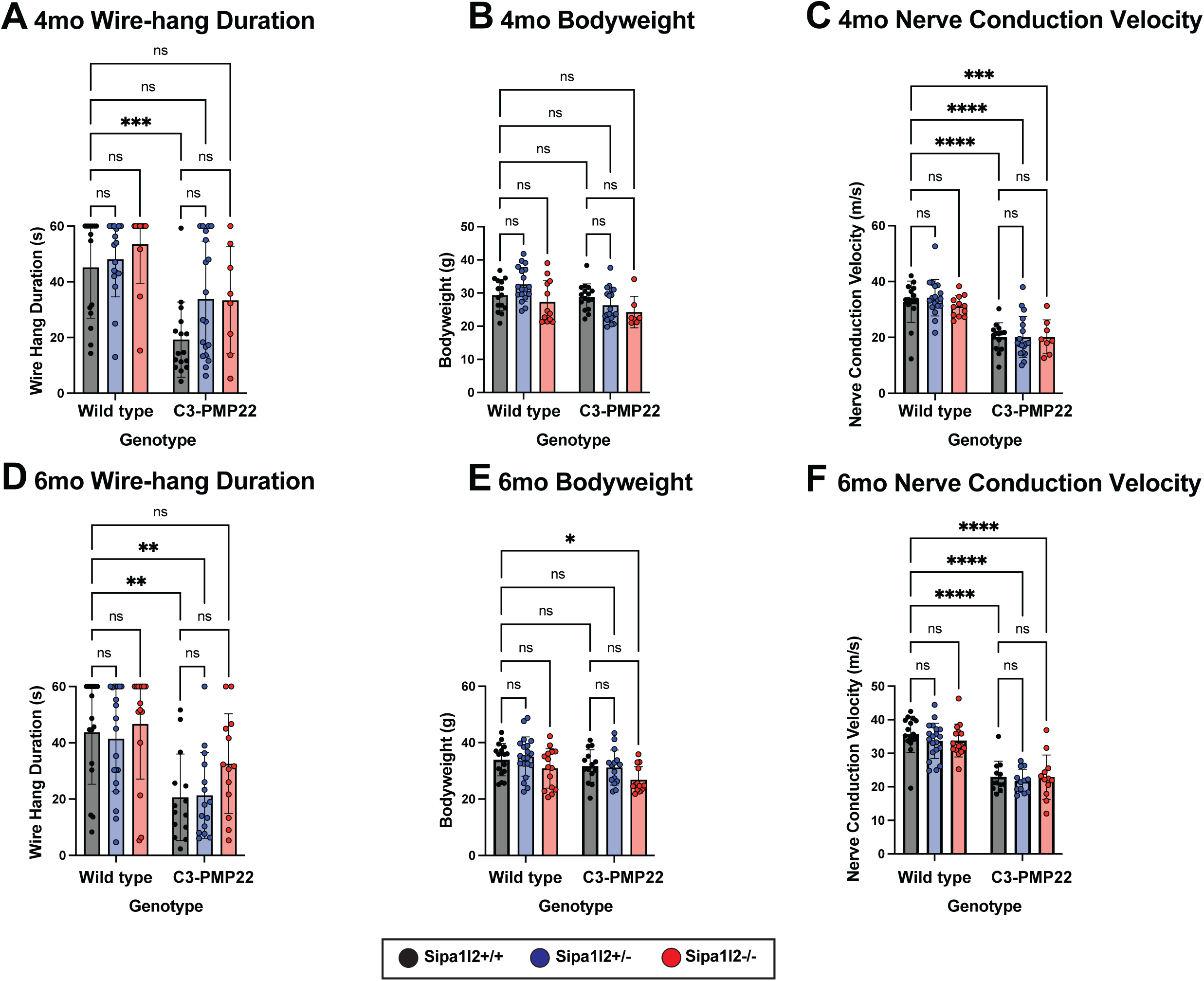
– Neuromuscular Phenotyping in 4mo and 6mo aged mice. A) Bar plot of wire-hang duration of 4mo old mice. Each point represents the average duration of 3 consecutive trials. C3-PMP22 mice have significantly reduced wire-hang duration (p=0.0007) compared to littermate wild-type mice. B) Bar plot of bodyweight of 4mo old mice. Each datapoint indicates one mouse. No significant differences are detected between genotype. C) Bar plot of sciatic nerve conduction velocity of 4mo old mice. Each data point represents one nerve conduction velocity measurement. Significant differences are detected between wild type and C3-PMP22 (p<0.0001), wild type and C3-PMP22::Sipa1l2+/-(p<0.0001), wild type and C3-PMP22::Sipa1l2-/-(p=0.0003). D) Bar plot of wire-hang duration of 6mo old mice. Each datapoint represents the average duration of three consecutive trials. C3-PMP22 and C3-PMP22::Sipa1l2+/-mice have significantly reduced wire-hang duration (p=0.0076 and p=0.0086, respectively) compared to wild-type littermates. E) Bar plot of bodyweight of 6mo mice. Each data point represents one mouse. Only C3-PMP22::Sipa1l2-/-mice differed significantly from wild types (p=0.012). F) Bar plot of sciatic nerve conduction velocity of 6mo old mice. Each datapoint presents one nerve conduction velocity measurement. Significant differences were detected between wild-type mice and C3-PMP22, C3-PMP22::Sipa1l2+/-, and C3-PMP22::Sipa1l2-/-(all p<0.0001). All comparisons are two-way ANOVA with either Tukey’s multiple comparisons test (wire-hang duration, nerve conduction velocity) or Dunnett’s multiple comparisons test (bodyweight). Error bars depict standard deviation.

### Histological reveals interactions between Sipa1l2 deletion and myelination

We collected motor and sensory branches of the femoral nerve at 6mo of age. Nerves were sectioned, stained with toluidine blue, and visualized at 40X by light microscopy. Genotypes expressing the C3-PMP22 transgene exhibit evident demyelination in motor branch of femoral nerve compared to those genotypes without the transgene, but this is effect is not clear in the sensory branch **(Figure 3)**. We counted the number of axons in both motor and sensory branches of the femoral nerve and identified no significant differences in the number of axons of femoral nerve motor branches between any genotypes **(Figure 4A)**. We detected significantly fewer axons in femoral nerve sensory branch for C3-PMP22 (p=0.0001) and C3-PMP22::Sipa1l2-/-(p<0.0001) mice, but interestingly, this reduction was not observed in C3-PMP22::Sipa1l2+/-mice **(Figure 4B)**. We next examined the ratio of totally demyelinated axons to total axons in both femoral nerve branches. All genotypes expressing the C3-PMP22 transgene exhibited significantly more demyelinated axons in the motor branch of femoral nerve **(Figure 4C)**. No difference between wild type mice and other genotypes in the percentage of demyelinated was detected in the sensory branch **(Figure 4D)**. It should be noted that any degree of myelination, even partial myelination, was scored as “myelinated.” Therefore, this analysis could miss effects on the degree of myelination in partially demyelinated axons, which were present in both motor and sensory branches.

**Figure 3.**
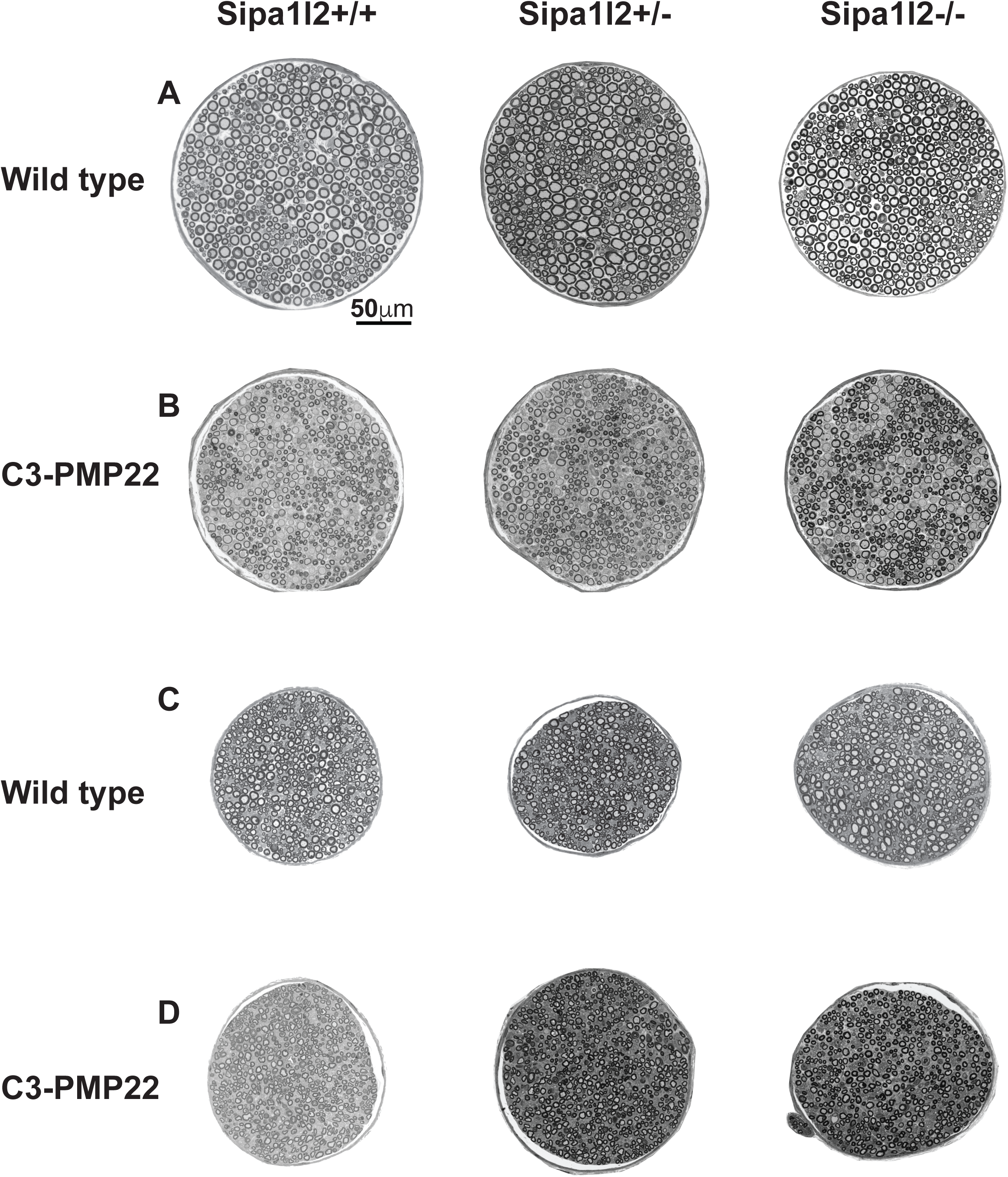
– Representative Histopathology of Femoral Nerve Branches. Representative images of toluidine blue stained sections from motor and sensory branches of femoral nerve visualized at 40X on a light microscope for all genotypes: A) motor branches of Sipa1l2 genotypes in a wild type background, B) motor branches of Sipa1l2 genotypes in the C3-PMP22 background, C) sensory branches of Sipa1l2 genotypes in a wild type background, D) sensory branches of Sipa1l2 genotypes in the C3-PMP22 background. Scale bar in panel A (50μm) applies to all images.

**Figure 4.**
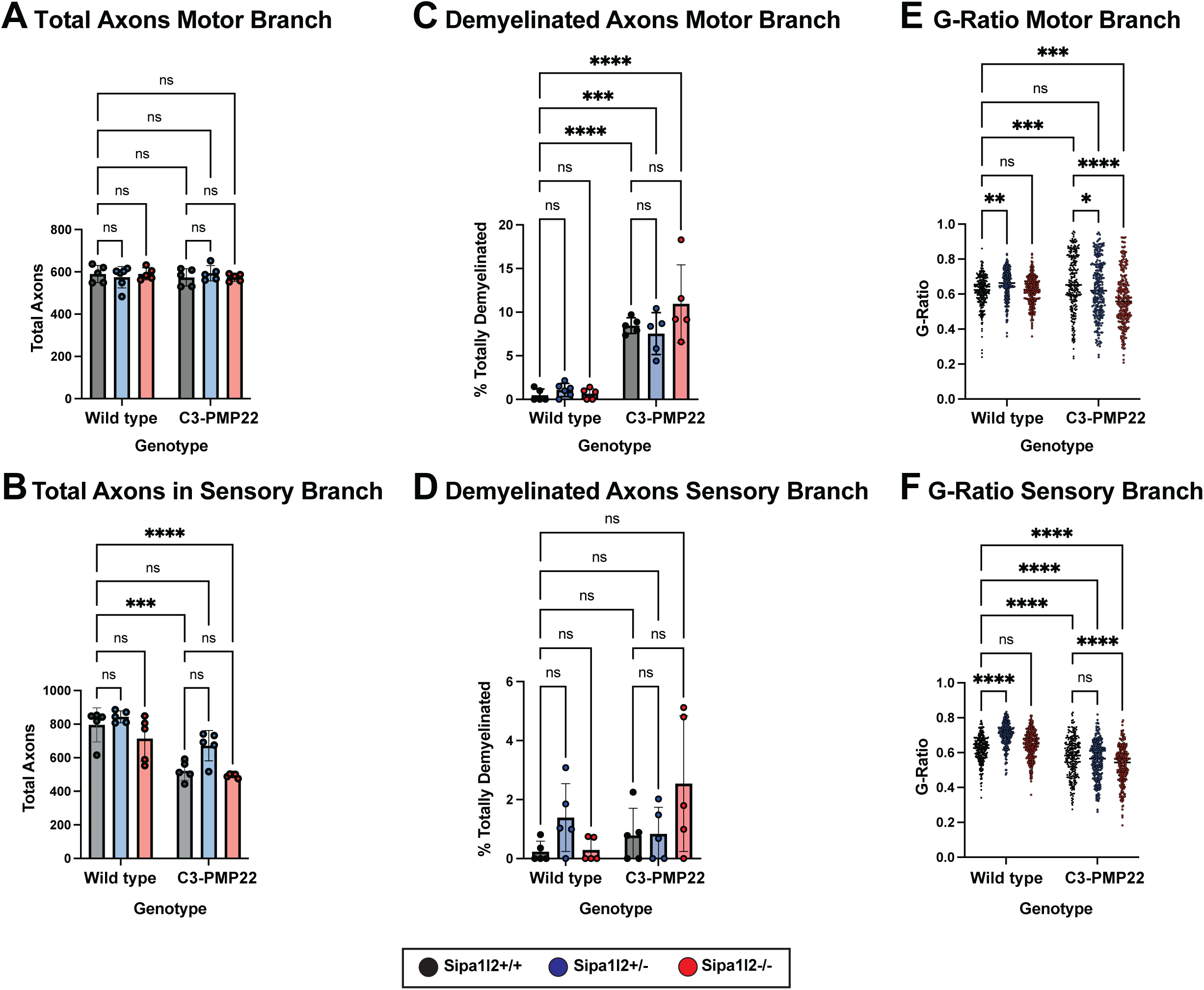
– Nerve Morphometrics from Femoral Nerve Motor and Sensory Branches. A) Bar plot of the total number of axons in the motor branch of the femoral nerve. No differences are detected between genotypes. B) Bar plot of the total number of axons in the sensory branch of the femoral nerve. Significantly reduced numbers of axons are detected between both C3-PMP22 (p=0.0001) and C3-PMP22::Sipa1l2-/-(p<0.0001). C) Bar plot depicting the percentage of totally demyelinated axons in the motor branch of the femoral nerve. Significant reduction is detected for all genotypes expressing the C3-PMP22 transgene (C3-PMP22 and C3-PMP22::Sipa1l2-/-p<0.0001, C3-PMP22::Sipa1l2+/-p=0.0002). D) Bar plot of the percentage of demyelinated axons in the sensory branch of the femoral nerve. No differences were detected. E) Dot plot depicting G-Ratio of axons from the motor branch of the femoral nerve. Significant differences are detected between wild type and Sipa1l2+/-(p=0.0063), wild type and C3-PMP22 (p=0.0005), wild type and C3-PMP22::Sipa1l2-/-(p=0.0009). F) Dot plot depicting G-ratio of axons from the sensory branch of the femoral nerve. Significant differences are detected between wild type and Sipa1l2+/-, C3-PMP22, C3-PMP22::Sipa1l2+/-, and C3-PMP22::Sipa1l2-/-(all p<0.0001). For axon counts and myelination ratios, each datapoint indicates one mouse. For g-ratios, 50 axons were randomly selected for quantification from five mice. All comparisons are two-way ANOVA with either Dunnett’s multiple comparisons test (axon counts) or Tukey’s multiple comparisons test (Percent demyelinated axons and G-ratios). Error bars depict standard deviation.

We next quantified g-ratios, defined as (axon diameter / axon + myelin diameter), for both branches of the femoral nerve. Thus, thinner myelin leads to a higher g-ratio, closer to a value of 1. In the motor branch, Sipa1l2+/-mice exhibit an increased g-ratio (p=0.0063), as do C3-PMP22 mice (p=0.0005), whereas C3-PMP22::Sipa1l2-/-exhibit a g-ratio significantly (p=0.0009) lower than wild types. Sipa1l2-/-and C3-PMP22::Sipa1l2+/-do not differ significantly from wild types **(Figure 4E)**. A similar increase above wild type is observed in the sensory branch of Sipa1l2+/-mice (p<0.0001). All genotypes carrying the C3-PMP22 transgene have significantly (p<0.0001) lower g-ratios in the sensory branch compared to wild type mice **(Figure 4F)**. When the relationship between axon diameter and myelin thickness is plotted, the effect is clearer. Mice without the C3-PMP22 transgene have a normal, upward slope, with larger axons having thicker myelin. C3-PMP22 mice have a downward slope, with larger axons having thinner myelin. G-ratio differences appear to be driven largely by an increase in myelination in the motor branch of the femoral nerve. Notably, while the downward slope is similar, C3-PMP22::Sipa1l2-/-mice have thicker myelin across the range of axon diameters (Prism Simple Linear Regression Y-Intercept ANCOVA p<0.0001). **(Supplemental Figure 1A)**. An influence of *Sipa1l2* deletion on myelin thickness is also detectable in the sensory branch of the femoral nerve; however, the effect is most obvious in Sipa1l2+/-mice on a wild type background, where heterozygous deletion reduces myelin thickness across all axon diameters **(Supplemental Figure 1B)**.

These results are complex, and g-ratios are difficult to interpret. In summary, we think there are several important takeaways. First, there is axon loss in the sensory branch of femoral nerve in genotypes with the C3-PMP22 transgene, except C3-PMP22::Sipa1l2+/-. Second, there are more totally demyelinated axons in all genotypes with the C3-PMP22 transgene in the motor branch of femoral nerve. Further, in motor branch, the C3-PMP22 transgene causes hypermyelination of small diameter axons and demyelination of large diameter axons, while Sipa1l2-/-deletion increases myelin thickness across all diameters in the C3-PMP22 background. In sensory branch, myelination trends are less obvious, but Sipa1l2+/-decreases myelin thickness in the wild-type background.

### RNASeq analysis of sciatic nerve implicates cholesterol biosynthesis

Library prep and sequencing was performed using sciatic nerves from wild type, Sipa1l2-/-, C3-PMP22, and C3-PMP22::Sipa1l2-/-mice because we presumed any interaction between SIPA1L2 and C3-PMP22 gene expression would be strongest in homozygous knockouts. Sciatic nerve was chosen because this tissue is rich in Schwann cells. Our differential expression analysis compared all experimental genotypes against wild types and used a false discovery rate cutoff of FDR<0.05, and we considered genes with an absolute Log_2_FC>1.5 to be differentially expressed **(Table S1)**. We identified 59 significantly differentially expressed genes in C3-PMP22 mice, 7 differentially expressed genes in Sipa1l2-/-mice, and 88 differentially expressed genes in C3-PMP22::Sipa1l2-/-mice **(Figure 5A)**. We also performed a targeted investigation of genes in the SOX10/EGR2 co-expression network and failed to identify any changes or differential expression in these genes in sciatic nerves at 6mo of age **(Supplemental Figure 2A)**. We also considered differentially expressed genes that were shared between multiple experimental genotypes. Only three genes overlapped between Sipa1l2-/-, C3-PMP22, and C3-PMP22::Sipa1l2-/-mice **(Supplemental Figure 2B)**. These genes are troponin I, ribosomal protein L34 pseudogene 1, and ribosomal protein S3A2. Their functions are related to calcium sensitivity in striated muscle, the large ribosomal subunit, and the small ribosomal subunit, respectively (STELZER *et al*. 2016).

**Figure 5.**
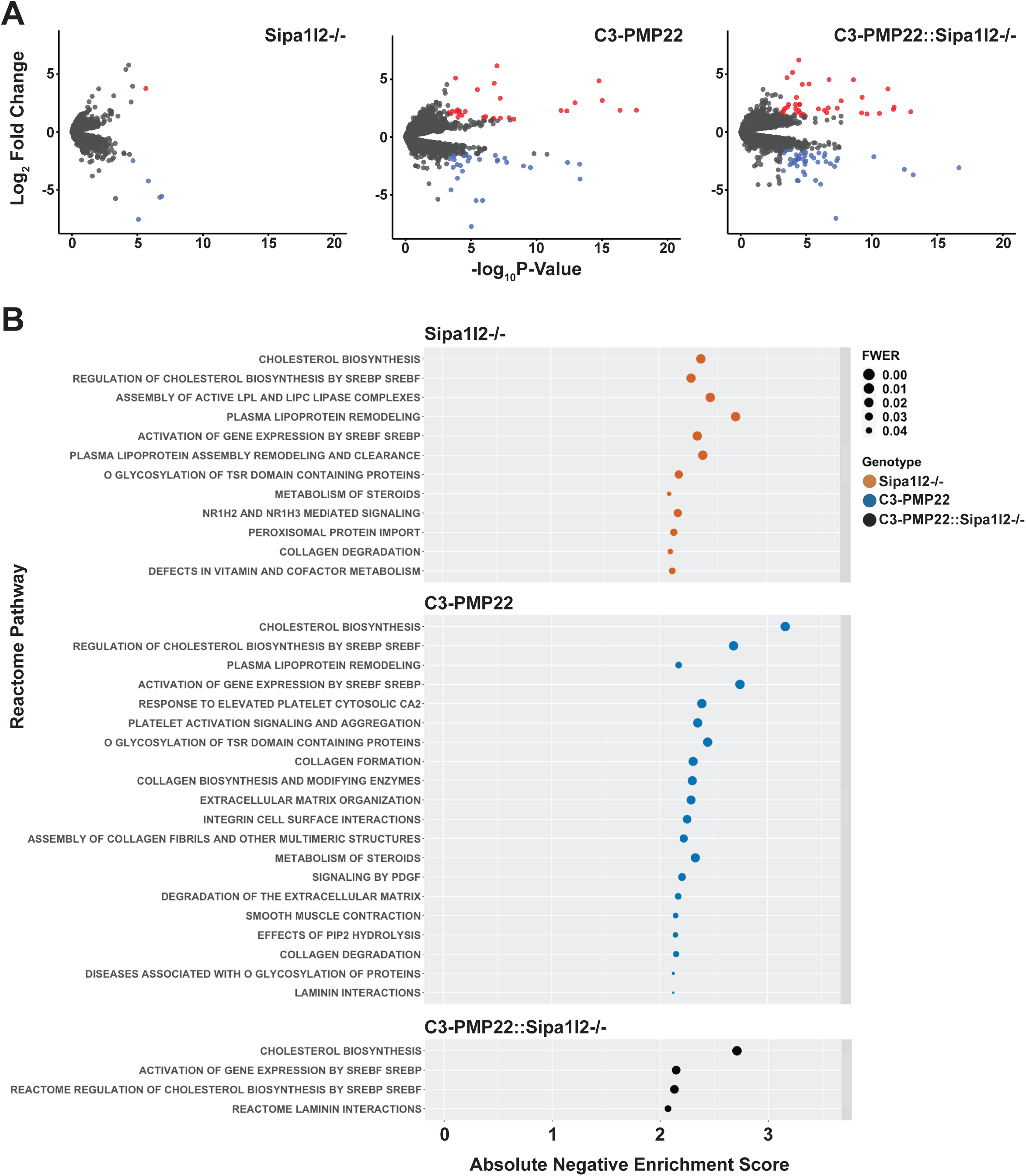
– Differential Expression and Gene Set Enrichment Analyses. A) Differentially expressed genes identified in comparisons between Sipa1l2-/-, C3-PMP22, and C3-PMP22::Sipa1l2-/-mice with wild type littermates were filtered by FDR<0.05 and absolute Log_2_FC>1.5. Gene lists were visualized by with volcano plots. Upregulated genes are colored red and downregulated genes are colored blue. Gray dots indicate genes that were filtered from analysis. We identified 7 DEGs for Sipa1l2-/-mice, 59 DEGs for C3-PMP22 mice, and 88 DEGs for C3-PMP22::Sipa1l2-/-mice. B) A dot plot depicting Reactome pathways identified for each experimental genotype based on TMM normalized counts data from RNASeq. Pathways with negative normalized enrichment scores are in descending order from the most significant family-wise error rate across genotypes, with an FWER < 0.05 cutoff. The x-axis reports the absolute value of the negative normalized enrichment score. In all cases, pathways are associated with a negative normalized enrichment score, or repressed. Genotypes are color coded.

We next performed a basic overrepresentation analysis using Mouse Mine. The gene list from C3-PMP22 mice was significantly associated with the Reactome pathway “striated muscle contraction” (Holm-Bonferroni p= 0.003211). Sipa1l2-/-mice gene lists were also associated with “striated muscle contraction” (HB p= 3.19e-7) and “muscle contraction” (HB p=1.60e-4). C3-PMP22::Sipa1l2-/-mice gene lists were not significantly associated with any Reactome pathways. Given the relatively low number of differentially expressed genes, and the paucity of overrepresented pathways associated with them, we decided to perform Gene Set Enrichment Analysis (GSEA) using TMM normalized counts values of all genes, which would better suited for detecting the coordinated effects of subtle expression changes in multiple genes, rather than using lists of differentially expressed genes.

GSEA Reactome pathways were filtered by family wise error rate (FWER<0.05) and ordered by absolute normalized enrichment score (NES) **(Table S2)**. Across mutant versus wild type contrasts, for all experimental groups, we found that pathways associated with upregulated genes in the experimental group (positive enrichment scores) tended to vary with enrichment statistic and metric for ranking genes, and the pathways themselves were diverse. Alternatively, those pathways with negative normalized enrichment score, which are frequently interpreted as downregulated or repressed in the experimental genotype, were robust across enrichment statistics, ranking metrics, and mutant genotypes. For this reason, we chose to focus our analysis on negative normalized enrichment scores. We found that cholesterol biosynthesis and pathways related to the regulatory activity of Sterol Response Element Binding Proteins / Factors (SREBP/SREBF) on cholesterol biosynthesis have large negative NES across Sipa1l2-/-, C3-PMP22, and C3-PMP22::Sipa1l2-/-mice **(Supplemental Figure 2C)**. Both Sipa1l2-/-and C3-PMP22 are associated with a variety of other significant pathways with lower magnitude NES, though C3-PMP22::Sipa1l2-/-is associated with fewer **(Figure 5B)**. Leading edge analysis of all genotypes identified lists of core enrichment genes primary responsible for the identification of cholesterol-related pathways **(Table S3) (Supplemental Figure 2D)**. Query of the ENCODE transcription-factor database through Network Analyst revealed that many of these leading edge genes are regulated by SREBP-1, providing a possible upstream regulator for the cholesterol-associated gene expression signatures.

## DISCUSSION

Here we report a newly CRISPR engineered strain of mice with a 5’-deletion in *Sipa1l2.* We did not detect any overt phenotypes, or neuromuscular phenotypes upon more detailed analysis, to be associated with the *Sipa1l2* deletion, but we did detect changes in g-ratio and myelin thickness in the motor and sensory branches of femoral nerve. Gene set enrichment analysis identified a group of Reactome pathways related to cholesterol biosynthesis and its regulation among downregulated pathways in the Sipa1l2-/-mice. We found several interactions between the *Sipa1l2* deletion and C3-PMP22 phenotypes. These include wire-hang duration, where *Sipa1l2* deletions promote endurance in mice overexpressing human *PMP22*. We also detected a slight reduction of bodyweight in transgenic mice homozygous for the *Sipa1l2* deletion. Notably, we did not detect an interaction between nerve conduction velocity deficits and the *Sipa1l2* deletion. Nerve conduction velocity decrements are an important feature of CMT1A, and demyelinating CMTs generally (BIROUK *et al*. 1997; KRAJEWSKI *et al*. 2000). We observed an interesting effect in nerve histology whereby heterozygous *Sipa1l2* deletions seem to prevent axon loss in the sensory branch of the femoral nerve. A heterozygous effect is also detected in motor branch of femoral nerve where wild type and C3-PMP22::Sipa1l2+/-mice do not differ significantly. Gene set enrichment analysis indicates repression of cholesterol biosynthesis across Sipa1l2-/-, C3-PMP22, and C3-PMP22::Sipa1l2-/-genotypes.

The effect of the *Sipa1l2-/-* deletion on wire-hang performance is perhaps most interesting because at both 4mo and 6mo timepoints C3-PMP22::Sipa1l2-/-mice are not distinguishable from wild types. C3-PMP22::Sipa1l2+/-mice, though not significantly different from wild type at 4mo, do show a significant decrease in endurance by 6mo. This establishes a dose effect, but that same dose effect is not present in our histopathology data. There, C3-PMP22::Sipa1l2+/-mice do not exhibit reduction in axon number in the sensory branch of femoral nerve, nor significant differences in motor branch g-ratio, compared to wild type mice while C3-PMP22::Sipa1l2-/-do. In the motor branch of femoral nerve, homozygous *Sipa1l2* deletion modestly increases myelin thickness. In this case, the increase in myelin thickness of transgenic mice with the homozygous *Sipa1l2* deletion may cause significantly different g-ratios from wild type while a more modest effect in transgenic mice with a heterozygous deletion in *Sipa1l2* maintain g-ratios closer to the wild type mean. Though the interaction between *Sipa1l2* deletion and myelination decrements in C3-PMP22 mice is interesting, it is not sufficient for functional rescue, as evidenced by our failure to detect differences in nerve conduction velocity. However, reduced nerve conduction velocity appears before other symptoms in *PMP22* duplication carriers, often in early childhood, which begs the question if complete rescue of myelination could improve conduction velocity (GARCIA *et al*. 1998; KRAJEWSKI *et al*. 2000; MANGANELLI *et al*. 2016).

The repression of gene sets related to cholesterol biosynthesis and its regulators such as sterol response element binding protein (SREBP) in *Sipa1l2-/-* mice compared to wild types is worth noting given the importance of local cholesterol synthesis in Schwann cells during myelin growth (JUREVICS AND MORELL 1994; GOODRUM *et al*. 2000; VERHEIJEN *et al*. 2009; KIM *et al*. 2018). There is evidence that coordinate action between SREBP and EGR2 regulates peripheral nerve myelination (LEBLANC *et al*. 2005; LI *et al*. 2013). Our finding that cholesterol biosynthesis genes are repressed without large changes in *Egr2* expression may suggest that either developmental changes in *Egr2* are sufficient to repress cholesterol biosynthesis in adult mice, or that SIPA1L2 performs a regulatory function in cholesterol biosynthesis independent of EGR2. Interestingly, rare variants in *SIPA1L2* have previously been associated with variable lipid response in humans, and rare variants in *SREBF1*, which encodes SREBP-1, have been implicated in Parkinson’s disease (DO *et al*. 2011; GENG *et al*. 2018). Though circumstantial, this may help explain why *SIPA1L2* variants are associated with Parkinson’s disease in only some GWAS studies (CHEN *et al*. 2016; SAFARALIZADEH *et al*. 2016; WANG *et al*. 2016; FOO *et al*. 2017; YANG *et al*. 2017; ZOU *et al*. 2018). It could be that *SIPA1L2* variants are only associated with Parkinson’s subtypes that exhibit altered cholesterol metabolism and the prevalence of individual patient subtypes differed between studies (MACIAS-GARCIA *et al*. 2021). Repression of cholesterol biosynthesis has also previously been reported in the C3-PMP22 mouse model, which we detect in our analysis as well (MICHAILIDOU *et al*. 2023). An increase in signal strength may explain why these same pathways are identified in C3-PMP22::Sipa1l2-/-mice, but without diverse lower enrichment score pathways that are also present in Sipa1l2-/-and C3-PMP22 mice. Alternatively, this may suggest that *Sipa1l2* deletion attenuates these pathway endophenotypes in C3-PMP22 mice.

Our *in vivo* results support the putative interaction between *SIPA1L2* and the severity of CMT1A phenotypes identified by patient GWAS (TAO *et al*. 2019a). Further, they may indicate an interaction through the highly relevant cholesterol biosynthesis pathway that exhibits transactivation with SOX10/EGR2 pathway components (LEBLANC *et al*. 2005). We did not detect SOX10/EGR2 pathway changes, and this may be due to developmental regulation or unknown activities of SIPA1L2. Though compelling, our neuromuscular phenotyping suggests that the effect of *Sipa1l2* deletion is relatively modest in the C3-PMP22 mouse model, and that effects on myelination do not abide by a straightforward dose effect. It is possible that a larger effect interaction may exist in rat models of CMT1A. The failure to improve nerve conduction velocity while also introducing a modest effect on bodyweight is disappointing from a functional therapeutic standpoint. We conclude that the *Sipa1l2* deletion modifies some CMT1A related phenotypes in mice, consistent with results from patient GWAS study, and that further study of *Sipa1l2* may help untangle the complex relationship between cholesterol biosynthesis and myelination in the peripheral nervous system during disease. However, the modest effect size and somewhat inconsistent gene dosage effects make modulation of SIPA1L2 a questionable therapeutic target for CMT1A.

## Supporting information

Differential Expression Analysis Results

Gene Set Enrichment Analysis Results

Leading Edge Analysis Results

## Acknowledgements

We gratefully acknowledge the contribution of Mouse Model Services and Genetic Engineering Technologies at the Jackson Laboratory for their assistance creating mice with the *Sipa1l2* deletion. We would like to thank Dr. Frank Baas for providing us with the C3-PMP22 mice. We also acknowledge the Electron Microscopy Core, particularly Pete Finger, at The Jackson Laboratory for expert assistance with the work described in this publication. We thank Genome Technology Services for their role in sequencing studies and Computational Sciences staff, particularly Grace Stafford, at The Jackson Laboratory for expert assistance with RNASeq analysis. Funding: This work was supported by NIH grants R21 NS116936, R24 NS098523, and R37 NS054154 to RWB. The Scientific Services at The Jackson Laboratory are supported by NCI grant CA34196. GCM was supported by T32GM132006.

## Author Contributions

GCM, TJH, and ALDT performed the experiments described and analyzed the results. IZ and SZ performed additional sequencing of CMT1A patient samples and analysis. The project was conceived and funded by RWB. GCM and RWB wrote the manuscript with input from all authors.

## Supplemental Figures

**Figure S1.**
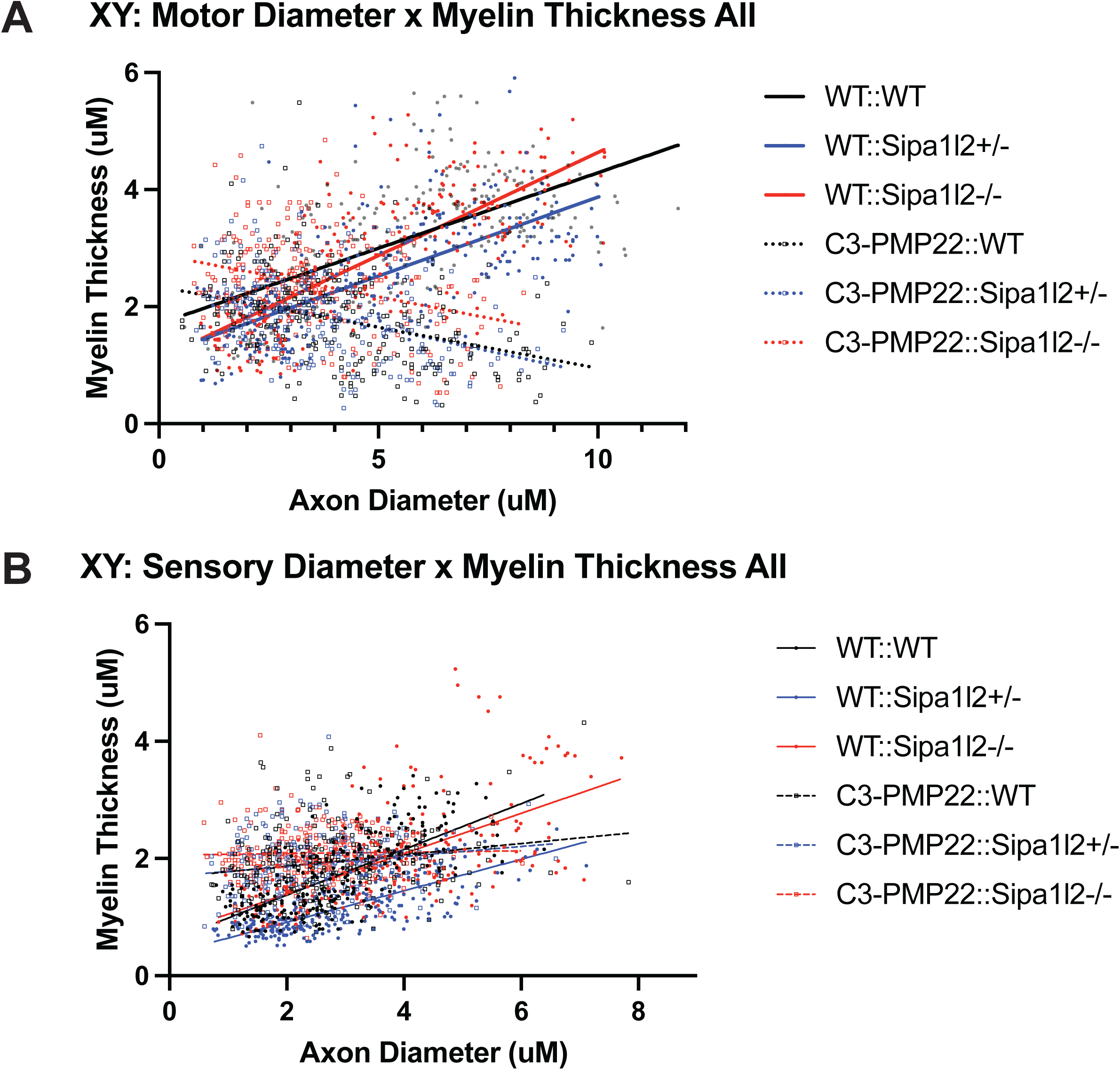
– Femoral Nerve Axon Diameter and Myelin Thickness XY Plots. A) An XY plot of axon diameter and myelin thickness for the motor branch of the femoral nerve. Slopes differ significantly (p<0.0001). B) An XY plot of axon diameter and myelin thickness for the sensory branch of the femoral nerve. Slopes differ significantly (p<0.0001). Each point indicates one axon. Fifty representative axons were randomly selected from five mice for quantification. Sipa1l2 deletion status is indicated with color. Dots and solid lines indicate mice without the C3-PMP22 transgene while hollow squares and dashed lines indicate transgenic C3-PMP22 mice. Simple linear regression in GraphPad Prism used to test for difference in slope of regression lines.

**Figure S2.**
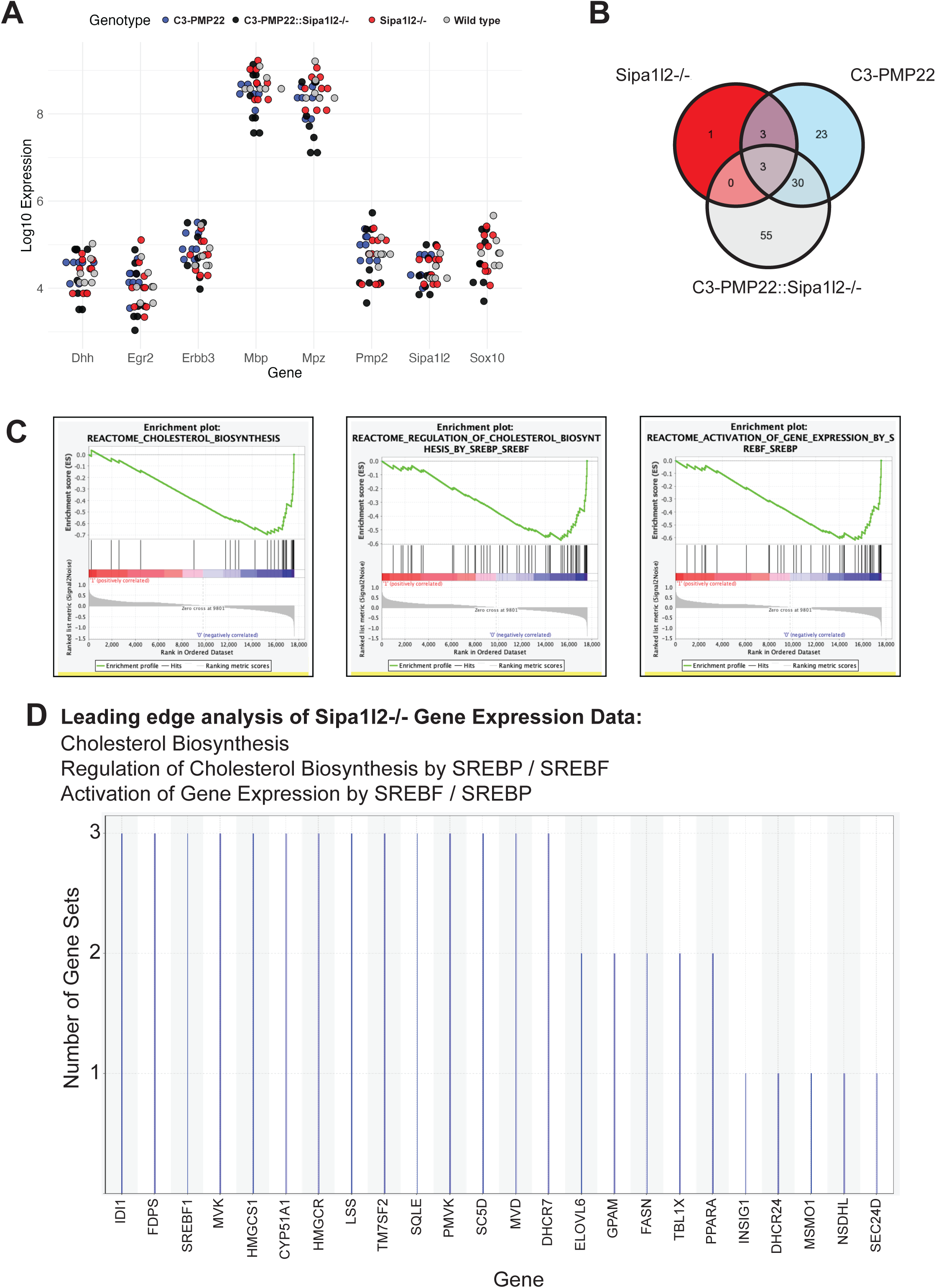
– Targeted Investigation of Gene Expression. A) Expression of genes associated with the SOX10/EGR2 co-expression network was investigated directly using TMM normalized counts from RNASeq. None of the selected SOX10/EGR2 genes were identified as differentially expressed between experimental genotypes. B) The overlapping differentially expressed genes between experimental genotypes were visualized using a Venn Diagram. Only three differentially expressed genes are shared between experimental genotypes. C) Representative gene set enrichment analysis enrichment (GSEA) plots for cholesterol-associated pathways in Sipa1l2-/-mice depicts negative enrichment or repression in mice with the deletion. D) Representative GSEA leading edge analysis identifies genes associated with cholesterol-associated pathways in Sipa1l2-/-mice.

## Supplemental Tables

**Table S1 – Differential Expression Analysis.** Contains the Ensembl identifiers, Log_2_Fold-Change, Log_2_CPM, p value, false discovery rate, and absolute Log_2_Fold-Change for all differentially expressed genes. Each experimental genotype comparison to wild type is contained in a different tab. Filtering can be adjusted to show genes that did not reach FDR or Log_2_FC cutoffs.

**Table S2 – Gene Set Enrichment Analysis.** Contains all Reactome pathways returned from gene set enrichment analysis. Gene set size, normalized enrichment score, and family wise error rate are reported for each pathway. Each experimental genotype is included in a different tab.

**Table S3 – Lead Edge Analysis.** Contains all genes returned from leading edge analysis of Reactome pathways related to “cholesterol” with FWER < 0.05. Each experimental genotype is provided in a column.

